# Hsp90 provides a platform for kinase dephosphorylation by PP5

**DOI:** 10.1101/2022.09.02.506407

**Authors:** Maru Jaime-Garza, Carlos Nowotny, Daniel Coutandin, Feng Wang, Mariano Tabios, David A. Agard

## Abstract

The Hsp90 molecular chaperone collaborates with the phosphorylated Cdc37 cochaperone to maintain kinase proteostasis through the folding and activation of its many client kinases. As with many kinases, the Hsp90 client kinase CRaf is activated by phosphorylation at specific regulatory sites. The cochaperone phosphatase PP5 dephosphorylates CRaf but also Cdc37 in an Hsp90-dependent manner. Although dephosphorylating Cdc37 has been proposed as a mechanism for releasing Hsp90-bound kinases, here we show that Hsp90 bound kinases sterically inhibit Cdc37 dephosphorylation indicating kinase release must occur before Cdc37 dephosphorylation. The cryo-EM structure of PP5 in complex with Hsp90:Cdc37:CRaf reveals how Hsp90 both activates PP5 and scaffolds its association with the bound CRaf to dephosphorylate a site at the C-terminus of the kinase domain. Thus, we directly show how Hsp90’s role in maintaining protein homeostasis goes beyond folding and activation to include post translationally modifying its client kinases.

## Introduction

Maintaining protein homeostasis is a critical function for all organisms and relies on a broad array of proteins including molecular chaperones^1^. The Hsp90 molecular chaperone is required for the folding and activation of over 10% of the human proteome^2,3^. Hsp90’s “client” proteins are enriched in signaling proteins such as protein kinases, transcription factors, and steroid hormone receptors. This leads Hsp90 to play an important role in organismal health and disease. Importantly, more than half of all human kinases depend on Hsp90 and its co-chaperone, Cdc37 for their folding and activation.^4^ The role of Hsp90 in kinase activation goes beyond folding and includes facilitating alterations in posttranslational modifications. Through its regulation of both kinase folding and kinase dephosphorylation, Hsp90 can modulate essential signaling pathways.

One such critical Hsp90-dependent pathway is the Ras-MAPK pathway involved in the regulation of cellular proliferation.^5,6^ When dysregulated, this pathway is often implicated in cellular malignancy.^7^ Raf kinases are part of this pathway, and act to propagate growth hormone signals from the membrane bound Ras GTPase to MEK and ERK kinases which can lead to signal amplification.^8^ Pathway activation requires Raf kinase dimerization which is mediated by phosphorylation of the Raf kinase acidic N-terminus.^9–11^ Thus, dephosphorylation of this acidic N-terminus may then lead to pathway inactivation.^12^

While Raf kinase activation has been extensively studied, Raf inactivation is less well understood. Importantly, for one member of the Raf family, CRaf (Raf-1), the Hsp90 cochaperone Protein Phosphatase 5 (PP5) has been directly implicated in its dephosphorylation and inactivation.^13^ More specifically, PP5 was shown to pulldown with CRaf and specifically dephosphorylate phospho-serine 338 (CRaf^pS338^) formed during Ras-MAPK pathway activation. Similarly, siRNA PP5 knockdown led to a specific increase in S338 phosphorylation. Based on these results, PP5 was hypothesized to play a key role in CRaf inactivation.

PP5 is a serine-threonine phosphatase from the PPP family which consists of a Tetratricopeptide (TPR) domain N-terminal to the catalytic phosphatase domain.^14,15^ The TPR domain sits directly atop the catalytic domain, sterically blocking substrate binding and access to the active site.^16,17^ Through predominantly hydrophobic interactions, an inhibitory αJ helix on the catalytic domain stabilizes the autoinhibited PP5 state through hydrophobic interactions with the TPR domain.^18^ Like other TPR-cochaperones, the PP5 TPR domain binds the Hsp90 C-terminal MEEVD tail, in this case leading to PP5 activation.^16,19^ A TPR domain mutation that blocks MEEVD binding abrogates the dephosphorylation of CRaf^pS338^ and inhibits PP5 coelution with CRaf kinase^13^. These results strongly suggest that Hsp90 plays a key role in CRaf dephosphorylation by controlling when and where PP5 becomes activated. In addition to CRaf, PP5 dephosphorylates numerous other Hsp90 clients, presumably while they are bound to Hsp90.^20–25^

Ppt1, the yeast homolog of PP5 has been shown to dephosphorylate yeast Hsp90 itself.^26^ This led Vaughan et al to hypothesize that PP5 might dephosphorylate the Hsp90 cochaperone Cdc37 which must be phosphorylated on S13 (Cdc37^pS13^) to function in kinase activation ^27–30^. Cdc37 is a kinase-specific Hsp90 cochaperone which binds and destabilizes kinase domains, enabling their recruitment into Hsp90:Cdc37:kinase complexes.^31^ Hsp90:Cdc37:kinase complexes purified from yeast, baculovirus or mammalian cells are invariably phosphorylated on Cdc37^S13. 29,32^ Various other groups have shown that the mutation of Cdc37^S13^ leads to a decrease in Hsp90 and kinase pulldowns.^28^ Finally, structural analysis of Hsp90:Cdc37:kinase complexes reveals that Cdc37^pS13^ is required to stabilize the Cdc37 N-terminal domain and to facilitate interactions with Hsp90.^33^

Without activation by Hsp90, PP5 by itself cannot dephosphorylate isolated Cdc37 or Cdc37 within a Cdc37:Cdk4 complex.^32^ PP5 can, however, dephosphorylate Cdc37 in the context of a purified Hsp90:Cdc37:Cdk4 complex whereas non-specific phosphatases cannot. This led to the hypothesis that PP5 must dephosphorylate Cdc37 while it is bound to an Hsp90:Cdc37:kinase complex and that it can thus facilitate kinase release from the Hsp90 complex.

To further understand the molecular mechanisms by which PP5 is activated and selects its target substrates, we solved the atomic resolution cryo-EM structure of an Hsp90:Cdc37:CRaf:PP5 complex and biochemically explored PP5-dependent dephosphorylation of both CRaf and Cdc37. Surprisingly, this revealed that while CRaf could be readily dephosphorylated, Cdc37 could only be dephosphorylated when activated by Hsp90 but in the absence of kinase. Through this work we propose a mechanism for PP5 activation and suggest that PP5 does not serve as a kinase release factor, but it may instead block rebinding to previously accessed Hsp90:Cdc37 complexes.

## Results

### Only kinase free Hsp90-Cdc37 complex can be dephosphorylated by PP5

In vivo, PP5 is reported to specifically dephosphorylate CRaf^pS338^ while leaving other essential CRaf phosphosites unaltered. In vitro, purified Hsp90:Cdc37:CRaf^304-648^ complexes are also dephosphorylated by PP5.^34^ Unless otherwise noted, purified phosphorylated CRaf^304-648^ complexes were used throughout our study and phosphorylation state was quantitatively assed using specific anti-phospho site antibodies directed against CRaf^pS338^(N-terminal to the kinase domain), as well as a control site C-terminal to the kinase domain (CRaf^pS621^)(Sup. Fig. 1)^13^. Additionally, anti-Cdc37^pS13^ antibodies were used to probe Cdc37 dephosphorylation.

PP5 activity was followed by incubating purified PP5 with purified Hsp90:Cdc37:CRaf (Fig. 1a, Sup. Fig. 2). CRaf ^pS338^ was promptly dephosphorylated. Surprisingly, the CRaf ^pS621^ control site was also dephosphorylated at about 40% the rate of CRaf ^pS338^. Contrary to our expectations, Cdc37^pS13^ was not measurably dephosphorylated upon PP5 addition. These results corroborate previous structural data showing that Cdc37^pS13^ is inaccessible within the complex, yet contradict previous experiments.^33^ A possible explanation is that differences in conformational homogeneity or partial complex dissociation might be required to enable Cdc37 dephosphorylation.

**Fig. 1:**
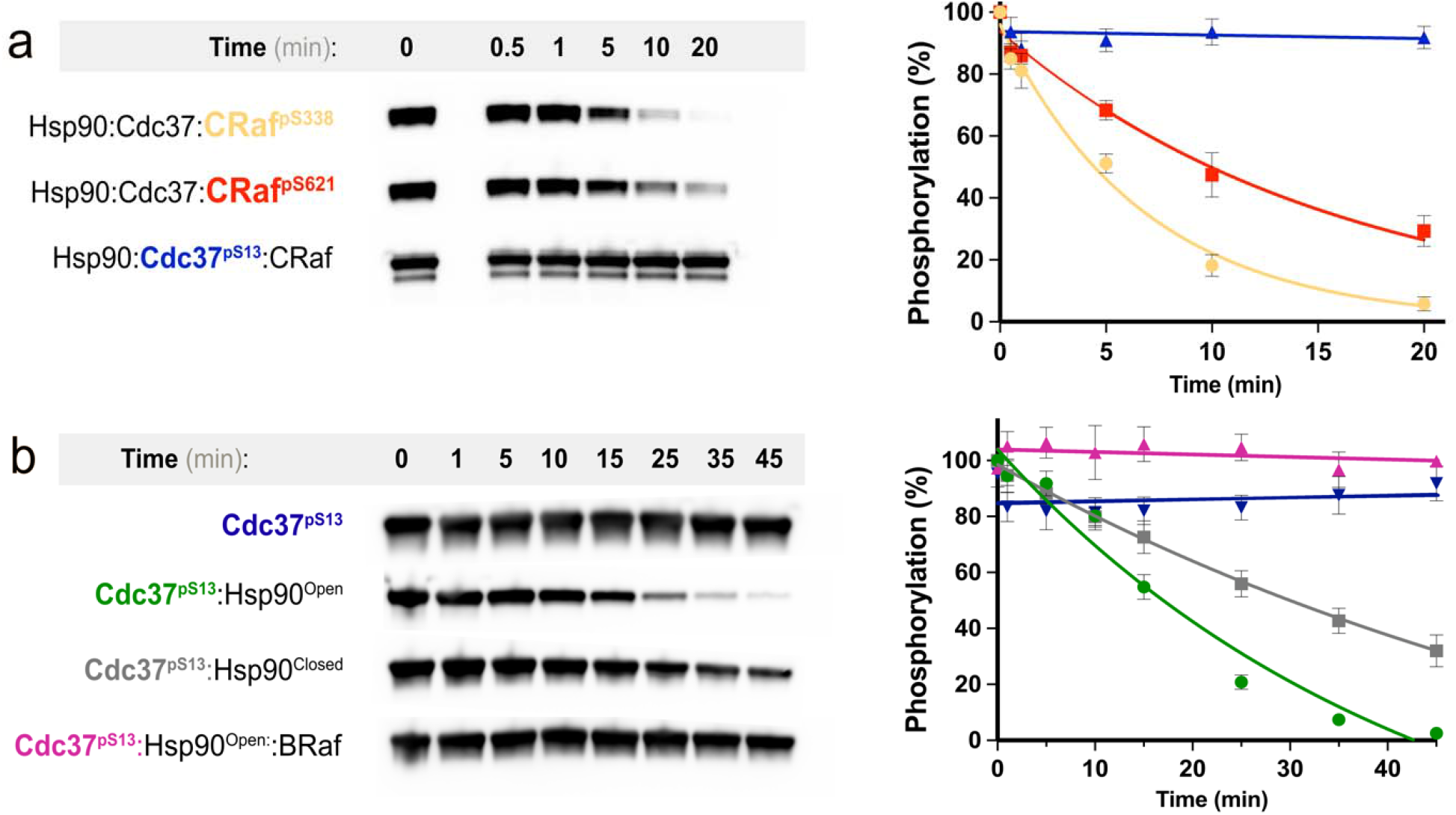
Kinase sterically blocks Cdc37 dephosphorylation. **a** Mammalian purified Hsp90-Cdc37-CRaf complex (1.5uM) was incubated with PP5 (75nM) at 25°C. The dephosphorylation of CRaf^pS338^, CRaf^pS621^ and Cdc37^pS13^ were assayed by phospho-specific blotting. CRaf^pS338^ was preferentially dephosphorylated (yellow, rate = 0.147 ± 0.012 min^-1^), CRaf^pS621^ was more slowly dephosphorylated (red, rate = 0.063 ± 0.006 min^-1^), while Cdc37^pS13^ was not dephosphorylated at all throughout this time course (blue, rate = 0.001 ±0.002 min^-1^). Both native and tagged Cdc37 are visible in the α-Cdc37^pS13^ blot. **b** Cdc37^pS13^ (3uM) was incubated with equimolar complex components (Hsp90^Open dimer^, Hsp90^Closed dimer^, BRaf) and PP5 (750nM) at 25°C. Cdc37^pS13^ dephosphorylation was assayed by α-Cdc37^pS13^ blotting. PP5 doesn’t dephosphorylate Cdc37 when Hsp90 is absent (blue, rate = 0.004 ± 0.37 min^-1^). PP5 dephosphorylates Cdc37^pS13^ bound to Hsp90^Open^ (green, rate = 0.024 ± 0.009) faster than when bound to Hsp90^Closed^ (gray, rate = 0.013 ± 0.01 min^-1^). There is no dephosphorylation with BRaf Kinase addition to Hsp90^Open^-Cdc37^pS13^ complex (magenta, rate = 0 min^-1^).

To probe the factors contributing to the inaccessibility of Cdc37^pS13^, we wanted to systematically explore the impact of Hsp90 nucleotide state and the role of the kinase in Cdc37^pS13^ dephosphorylation. As shown by cryo-EM, natively isolated Hsp90:Cdc37:kinase complexes have a closed Hsp90 with an inaccessible Cdc37^pS13^.^33^ In principle, by leaving out the nucleotide in an in vitro reconstitution, it should be possible to form Hsp90 open-state complexes. Unfortunately, it has not yet been possible to reconstitute CRaf assembly into an Hsp90 complex in vitro. However, it is possible to assemble Hsp90:kinase domain complexes using the heavily mutated and solubilized BRaf kinase domain.^35,36^ Hsp90, Cdc37, and BRaf were expressed and purified from E. coli, and Cdc37 was phosphorylated by CK2 before a final purification. We reconstituted an Hsp90:Cdc37:BRaf complex by mixing and incubating for 30min at 4°C. Dephosphorylation of Cdc37^pS13^ upon addition of PP5 was again quantitated (Fig. 1b). In support of our results, PP5 was also unable to dephosphorylate Cdc37^pS13^ reconstituted into Hsp90^open^ Hsp90:Cdc37:BRaf complexes.

To test if BRaf might sterically occlude PP5 from acting on Cdc37, we carried out the phosphatase reaction without the kinase present. Notably, Cdc37^pS13^ was rapidly dephosphorylated. We reasoned that if accessibility were the main factor limiting Cdc37^pS13^ dephosphorylation, then closing Hsp90 without BRaf present might hinder PP5 access. Hsp90 was incubated with AMPPNP and allowed to shift to a closed conformation after which Cdc37^pS13^ was added and dephosphorylation was measured.^37^ Interestingly, Hsp90^closed^:Cdc37^pS13^ was dephosphorylated at about half the rate as Hsp90^open^:Cdc37^pS13^, suggesting that while the Hsp90 conformational state contributes to the occlusion of Cdc37^pS13^, activity is dominated by the presence or absence of kinase.

### PP5 becomes activated and uses Hsp90 as a dephosphorylation scaffold

Since there was strong PP5 activity on CRaf, we focused on determining the atomic structure of activated PP5 using an Hsp90:Cdc37:CRaf:PP5 complex. Following from previous cryo-EM studies, we natively isolated stable Hsp90^closed^ :Cdc37:CRaf complexes from yeast using just the CRaf kinase domain (CRaf^KD^, residues 336-648). An inactive PP5 mutant PP5^H304A^ was used to stabilize any substrate bound PP5 complex. To keep the complex intact through vitrification and ensure protection from dissociation at the air-water interface, we chemically crosslinked the complex and froze it on grids covered with a monolayer of graphene oxide derivatized with amino-PEG groups (Sup. Fig. 3).^38,39^ A large single particle data set was collected and processed using Cryosparc and Relion.

Despite having a biochemically homogeneous crosslinked sample, data processing indicated a conformationally heterogeneous complex requiring significant 3D classification and refinement yielding a 3.3Å map (Fig. 2a). In this map, Hsp90 sidechains could be readily interpreted and more detail for Cdc37 and the kinase was observed than in our previous, lower resolution Hsp90:Cdc37:Cdk4 kinase complex structure.^33^ As expected from the Cdk4 kinase complex, the CRaf^KD^ kinase domain was split into its distinct lobes, threaded through the Hsp90 lumen and stabilized by Cdc37. The new map shows a closed Hsp90, with the CRaf^KD^ C-lobe extending from the Hsp90 lumen on one side as well as the prominent Cdc37 N-terminal coiled-coil projecting away from the complex surface. In the Cdk4 complex structure, only very low-resolution density could be seen for the Cdk4 N-lobe or the Cdc37 middle domain (Cdc37^MD^), and these features were mutually exclusive. By contrast, in the refined CRaf map we could directly observe the β4-strand from the kinase N-lobe bound to a groove in the Cdc37^MD^ (to be reported in more detail separately).

**Fig. 2:**
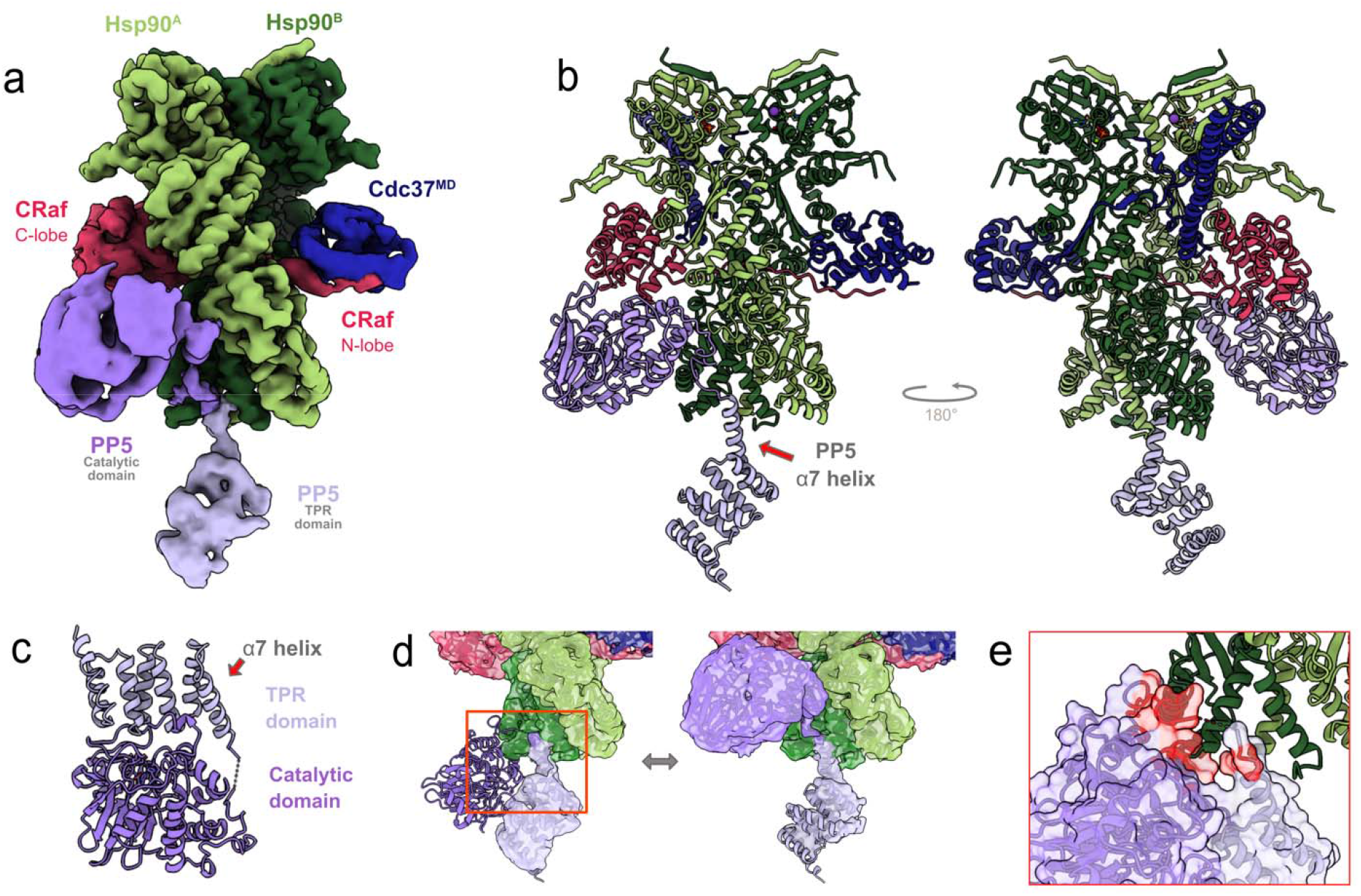
Hsp90 activates PP5, acting as a phosphatase scaffold. **a** CryoEM composite map and Hsp90:Cdc37:CRaf:PP5 complex model (**b**) shows unfolded CRaf kinase (magenta) threaded through the closed Hsp90 dimer lumen (green), and embraced by Cdc37 (blue). PP5 is in an active conformation as its TPR domain (lavender) interacts with the Hsp90 CTDs, while the PP5 catalytic domain (purple) interacts with Hsp90^MD^ and the CRaf C-lobe. **c**. In the PP5 crystal structure (PDB: 1wao) the PP5 active site is occluded when not bound to Hsp90. **d** Overlaying the autoinhibited PP5 structure PP5 with ours reveals steric clashes (**e**, in red) with Hsp90 that would necessitate separation of PP5 domains and drive PP5 activation.

Beyond these new features within the kinase complex, two new densities were visible near both Hsp90 C terminal domains (Hsp90^CTD^) and the Hsp90 middle domain (Hsp90^MD^) of protomer A. Focused local 3D classification around the newfound densities provided higher resolution views resulting in the composite map shown in Fig. 2a (Sup. Fig. 4).

A complete atomic model (Fig. 2b) constructed using the consensus and composite maps clearly reveals a PP5 conformation quite distinct from that found in the autoinhibited crystal structure (Fig. 2c).^16,40^ Instead, the PP5 TPR domain α7 helix is nestled into a groove between the two CTDs of Hsp90, making an end-on interaction with Hsp90 (Fig 2d). While the same Hsp90 groove is used for binding the TPR containing Fkbp51 cochaperone (Sup. Fig. 5)^37^, the Fkbp51 α7 helix lies perpendicular to the Hsp90 dimer, rotated almost 90° from the PP5 α7 helix orientation. In the complex, the two very separated PP5 domains are connected by an extended linker that is sufficiently ordered to be visible as it traverses a path along the surface of the Hsp90^CTD^s. Overlaying the TPR domains from our structure and the autoinhibited state reveals that the catalytic domain would severely clash with the Hsp90^CTD^ (Fig. 2e), necessitating TPR domain-catalytic domain separation. These interactions directly facilitate PP5 activation.

**Fig. 3:**
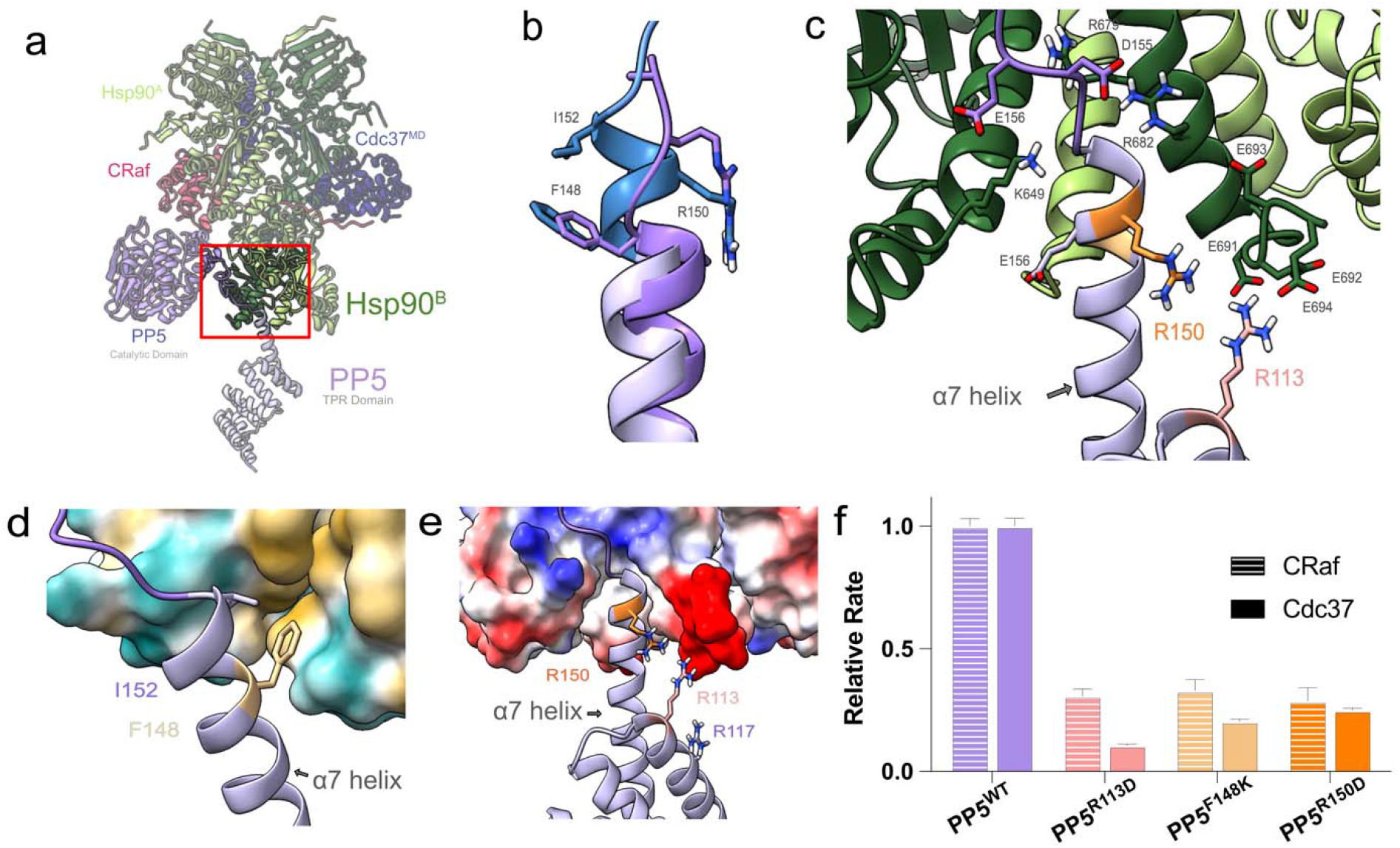
Novel TPR a7 helix - Hsp90 C-terminal domain interaction mode. **a The** PP5 TPR domain interacts with the amphipathic Hsp90 dimer C-terminal groove. **b** Upon Hsp90 interaction, the PP5 α7 helix elongates (lavender + blue) as compared to the inhibited PP5 crystal structure (purple, PDB: 1wao) **c** Electrostatic interactions between the PP5 linker and Hsp90 can be seen in more detail. **d** Hydrophobic residues PP5^F148^ and PP5^I152^ bind the Hsp90 hydrophobic CTD groove, while (**e**) PP5^R150^, PP5^R113^ electrostatically interact with the Hsp90 acidic tail. **f** Mutations to key TPR residues significantly decrease PP5 dephosphorylation of both Cdc37 and CRaf^pS338^ substrates (PP5^WT^ rate = 0.182±0.018 min^-1^, PP5^R113D^ = 0.056±0.017 min^-1^, PP5^F148K^ = 0.060±0.027 min^-1^, PP5^R150D^±0.033 min^-1^). The data was normalized to a PP5 free control, and the dephosphorylation rates were normalized to WT activity. The differences between PP5 mutants and PP5^WT^ were proven significant with an Ordinary one-way ANOVA (CRaf: n=3, F = 72.12, p < 0.0001, DF = 43; Cdc37: n^mutants^=3, n^WT^=4 F=411.9, p<0.0001, DF=60) with a Dunnett test for multiple hypothesis testing.

**Fig. 4:**
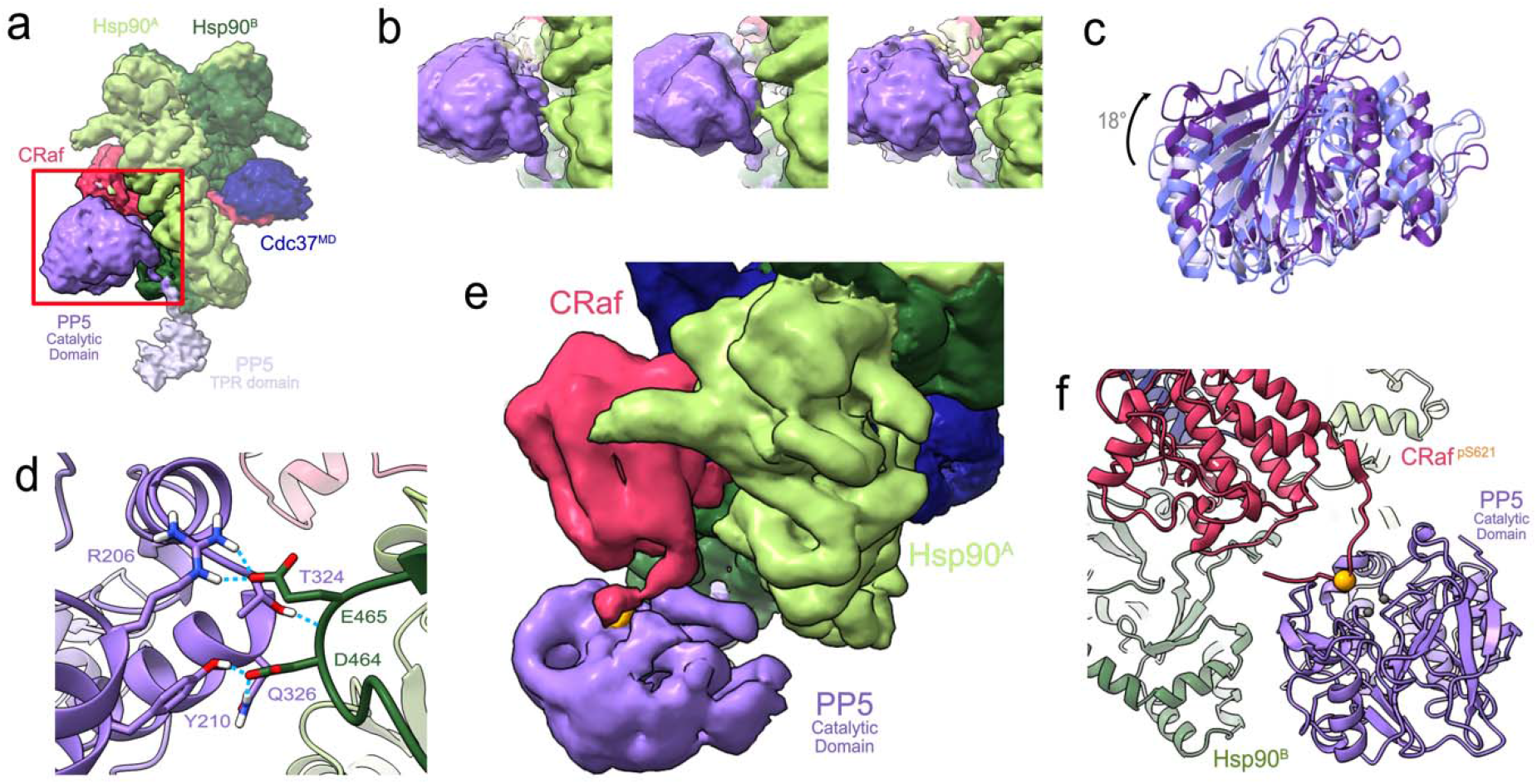
PPG’s catalytic domain interacts with Hsp90^MD^ and dephosphorylates CRaf^pS621^. **a** The PP5 catalytic domain binds to Hsp90^MD^ and the CRaf kinase C-lobe. **b** PP5 flexibly contacts Hsp90 allowing it to pivot about the interface. **c** Focused classification, alignment by Hsp90 density, and rigid docking demonstrates multiple PP5 conformations involved in the heterogeneous interaction with Hsp90^MD^. **d** A schematic of the Hsp90^MD^-PP5 interaction demonstrates potential electrostatic interactions and hydrogen bonding between an acidic loop on Hsp90 and the PP5 catalytic domain. **e** Focused classification of the PP5-CRaf interface shows low resolution density for the CRaf C-terminal tail directed towards the PP5 active site. **f** A model of the CRaf^pS621^ (orange) phosphorylated tail extending outward from the CRaf C-lobe was built through comparison with a crystal structure of substrate bound PP5 (PDB: 5hpe). The model shows an ideal length for CRaf^pS621^ substrate positioning in the PP5 active site.

**Fig. 5:**
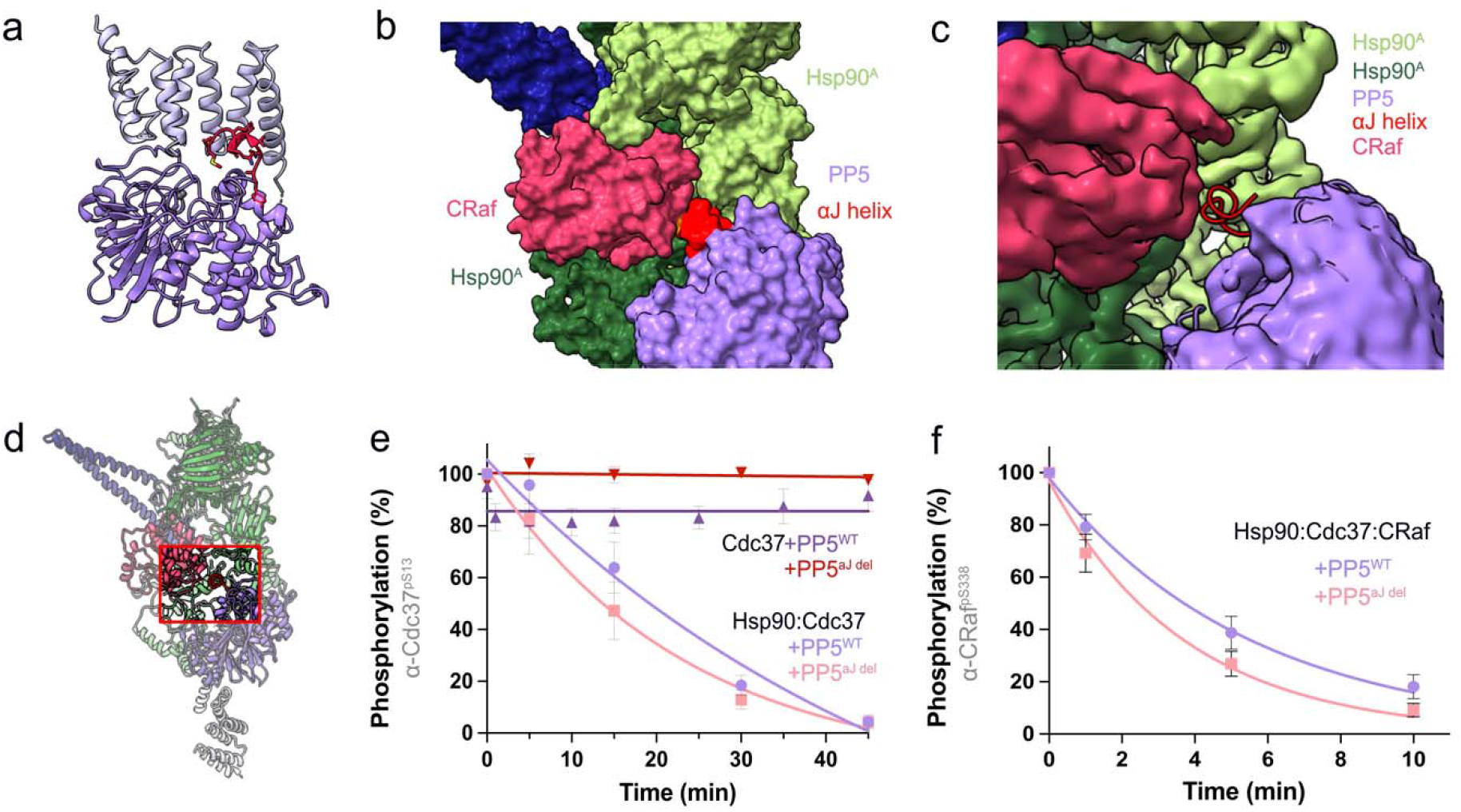
Deletion of PP5’s αJ helix increases rate of dephosphorylation. **a** In its inhibited conformation, the PP5 αJ helix (red) lies at the interface between the PP5 catalytic and TPR domains. **b** Without rearrangement, the inhibited conformation of the PP5 αJ helix would sterically clash with CRaf upon the activation of PP5 by Hsp90. **c**,**d** No αJ helix density is seen where expected in focused 3D classified catalytic domain maps, suggesting that the αJ helix is significantly disordered. **e** PP5 ^αJ del^ does not dephosphorylate Cdc37 without Hsp90 (red, rate = 0.0 ± 0.001, but PP5^αJ del^ dephosphorylates Cdc37 while bound to Hsp90 (pink, PP5^αJdel^ rate = 0.058 ± 0.009 min^-1^) at a higher rate than PP5^WT^ (lavender, PP5^WT^ rate = 0.047 ± 0.007 min^-1^). **f** PP5 ^αJdel^ leads to a slight increase in CRaf^pS338^ dephosphorylation when compared to PP5^WT^ (pink, PP5^αJdel^ rate = 0.27 ± 0.03 min^-1^ ; lavender, PP5^WT^ rate = 0.182 ± 0.018 min^-1^). The differences between PP5 ^αJ del^ and PP5^WT^ dephosphorylation rates were proven significant with unpaired t-tests (Cdc37^pS13^: two-tailed p = 0.0004, t =4.01, df=31 ; CRaf^pS338^: two-tailed p < 0.0001, t=7.823, df=20).

Surprisingly, the PP5 catalytic domain was not located near the readily dephosphorylated CRaf^pS338^ but was instead located on the other side of the Hsp90 dimer with the PP5 active site facing the CRaf C-lobe. Although CRaf residues beyond CRaf^616^ are disordered in our map, an approximate mainchain path connecting to the active site can be seen (Fig. 4e,f). This would place CRaf^pS621^ very close to the active site making it readily accessible for dephosphorylation. Combined with our dephosphorylation data (Fig 1a), this suggests that PP5 can act on CRaf^pS621^ while bound to Hsp90 as seen here.

### PP5’s TPR domain interacts extensively with Hsp90’s C-terminal domains via an extended helix

In its active state, the PP5 α7 helix binds in the amphipathic grove formed by CTD helices from each Hsp90 protomer (Fig. 3a). Binding at this interface stabilizes and elongates the C-terminal end of the α7 helix by a turn (Fig. 3b). Interactions towards the Hsp90 surface are primarily electrostatic while those deep in the CTD groove are hydrophobic (Fig. 3c).

The PP5 linker connecting the catalytic domain to the α7 helix has conserved acidic residues (PP5^D155^ and PP5^E156^) that lie in close proximity to Hsp90^R679^ and Hsp90^R682^ (Fig. 3c), stabilizing the α7 helix within the Hsp90 amphipathic groove. Here, the PP5^F148^ and PP5^I152^ residues which were solvent exposed and disordered in the PP5 autoinhibited state, are stabilized by Hsp90 hydrophobic residues (Hsp90^A^: L638, L654, L657 and Hsp90^B^: I680, I684, M683, L686) (Fig. 3d, Sup. Fig. 7).

Density corresponding to the Hsp90 tail can be seen extending beyond the last Hsp90^CTD^ α–helix towards the PP5 TPR domain. While poorly resolved, the charge complementarity between the Hsp90 tail (Hsp90^D691-E694^) and basic residues on the PP5 TPR domain (PP5^R150^, PP5^R113^, PP5^R117^) suggest that Hsp90 and PP5 interact beyond the α7 helix (Fig. 3e, Sup. Fig. 6a). This electrostatic interaction may guide the Hsp90 tail towards the PP5 MEEVD binding site, for which density is visible in our map (Sup. Fig. 6c, d).

The “entrance” for the TPR α7 helix on the Hsp90 CTD pyramidal groove measures approximately 13Å wide when Hsp90 is in a closed position, but contracts by almost 3Å when Hsp90 is in the Hsp90 partially open conformation found in the Hsp90:Hsp70:Hop:GR client loading state (Sup. Fig. 8).^41^ This suggests that PP5 binding might be substantially weaker in the fully open apo state or the partially open client loading state.

To determine the functional significance of the novel PP5-Hsp90^CTD^ interactions observed here we mutated three key residues (PP5^R113D^, PP5^F148K^, PP5^R150D^) and assessed the impact on both CRaf ^pS338^ and Hsp90^open^:Cdc37^pS13^ dephosphorylation (Fig. 3f). All three mutations led to ∼3-fold reduction in dephosphorylation rate for CRaf ^pS338^ and a somewhat greater impairment on Hsp90^open^:Cdc37^pS13^. This confirms the broad importance of the PP5 TPR-Hsp90 interactions for both substrates. The larger impact of these mutations on Cdc37 dephosphorylation may be due to the reduced size of the pyramidal pocket on more open Hsp90 states, further lowering affinity.

### Hsp90 positions the PP5 catalytic domain to efficiently dephosphorylate CRaf ^pS621^

Although the PP5 catalytic domain is clearly observed interacting with Hsp90^MD^, it is not rigidly anchored (Fig. 4a,b). Instead, 3D local classification reveals that it can pivot up to ∼18° about the Hsp90 interaction interface (Fig. 4c). The major area of interaction is between PP5 residues R206, Y210, T324, Q326 and an Hsp90^MD^ acidic loop (Hsp90^D464^ and Hsp90^E465^) (Fig. 4d). In keeping with the observed flexibility, we estimate that only ∼330 Å^2^ are buried in this interface, suggesting that it is a relatively weak interaction.

The PP5 active site faces towards the CRaf C-lobe. Focused classification on the interface between the two domains reveals continuous density extending from the last well resolved helical residue in the CRaf C-terminus (CRaf^611^) to within 8Å of the PP5 active site (Fig. 4e). While the density was too poor to accurately model de novo, a plausible model (Fig. 4f) was constructed based on the covalently bound PP5-substrate crystal structure (pdb: 5hpe). This provides a molecular explanation for the location of the PP5 catalytic domain and our unexpected observation that PP5 potently dephosphorylates CRaf^pS621^ in vitro.

### PP5^αJ helix^ is displaced upon interaction with Hsp90:Cdc37:CRaf complex

The PP5 C-terminal αJ helix (red) makes stabilizing interdomain interactions in the autoinhibited PP5 crystal structure (Fig. 5a). If the αJ helix were to remain in the same conformation upon PP5 activation (Fig. 5d), it would sterically clash with the CRaf C-lobe in the catalytically active domain-separated state (Fig. 5b). Thus, some rearrangement of the αJ helix is required. No density is observed for the αJ helix in our maps (Fig. 5c), suggesting that it becomes highly disordered upon PP5 activation.

From these observations we would expect removal of the αJ helix to be catalytically activating. To test this, we truncated the last 10 PP5 residues (PP5^αJ del^). As shown in Fig 5e, any resultant destabilization of the autoinhibited state was clearly insufficient to activate PP5 in the absence of Hsp90 (Fig. 5e). However, PP5^αJdel^ did lead to a slight increase in the rate of CRaf^pS338^ and Cdc37^pS13^ dephosphorylation in the context of the relevant Hsp90 complexes (Fig. 5f). Together these results suggest that the αJ helix stabilizes the inactive PP5 conformation, limiting PP5’s ability to bind Hsp90 and thus acts to decrease the rate of dephosphorylation.

## Discussion

PP5 is a unique member of the PPP family of phosphatases in that it contains an autoinhibitory TPR domain, and does not require coupling to separate cofactors for regulation.^42^ Through its interaction with Hsp90, PP5 benefits from Hsp90 client unfolding and holding, exposing phosphorylation sites that are otherwise inaccessible or too flexible for other phosphatases.

PP5 acts as a negative regulator for numerous Hsp90 clients and may reset Hsp90 clients to a basal phosphorylation state. This is the case with Hsp90 kinase clients such ASK1 kinase in oxidative stress, ATM kinase in DNA double strand break stress, and Chk1 kinase in UV induced stress.^20,22,25^ Not limited to kinases, PP5 also inactivates other Hsp90 clients such as Hsf1, p53 and the glucocorticoid receptor in an Hsp90 dependent manner.^21,43,44^

We set out to understand how Hsp90 activates PP5, and how Hsp90 can help position PP5 for efficient client dephosphorylation. Our Hsp90:Cdc37:CRaf^KD^:PP5 complex structure illustrates the molecular mechanism underlying both activities. The PP5 TPR and catalytic domains are separately bound to Hsp90, activating PP5. The PP5 α7 helix in the TPR domain makes unique interactions with the Hsp90^CTD^ groove different from other TPR containing Hsp90 cochaperones. The PP5 catalytic domain is positioned by the Hsp90^MD^ to interact with the CRaf^C-lobe^, where the CRaf^pS621^ phospho-site is dephosphorylated by PP5. While the shorter CRaf^KD^ may preclude the visualization of PP5:CRaf^N-lobe^ interactions by cryo-EM, we hypothesize that PP5 may adopt a mirrored conformation to that seen in our structure to dephosphorylate CRaf^pS338^ (Fig. 6a, Extended Fig. 10).

**Fig. 6:**
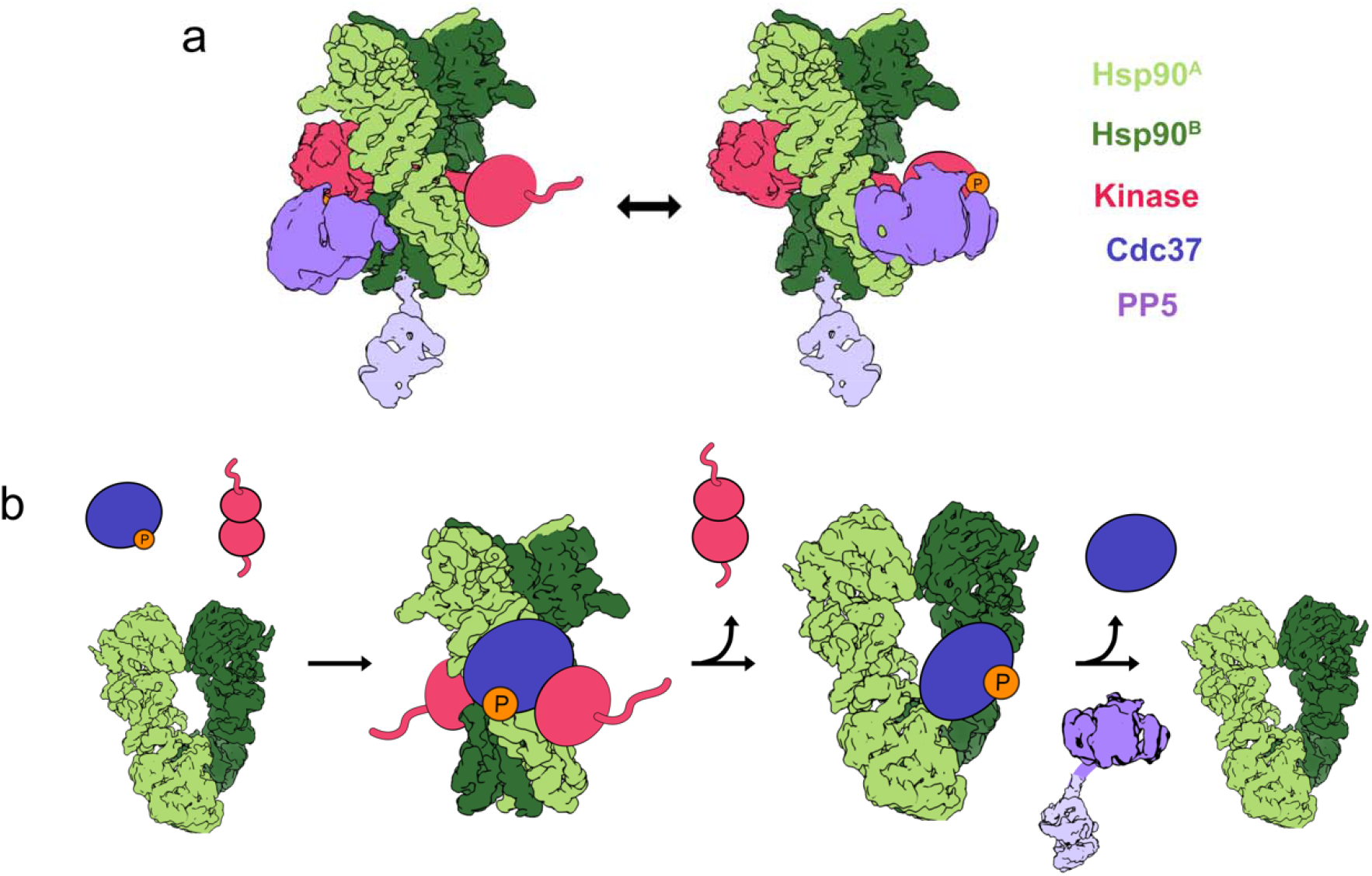
Mechanisms of kinase and Cdc37 dephosphorylation by PP5. Models illustrating the ability of Hsp90-bound PP5 to dephosphorylate residues on either side of Hsp90-bound clients and the role of PP5 in dephosphorylating Cdc37. **a** The PP5 TPR domain binds the amphipathic groove between the Hsp90^CTD^s, which activates the PP5 catalytic domain and allows it to use Hsp90 as a scaffold for client dephosphorylation on either side of Hsp90. **b** Steric blocking of Cdc37^pS13^ dephosphorylation by kinase clients suggests that Cdc37 can only be dephosphorylated once the kinase has left the Hsp90 complex. In our model, Cdc37^pS13^ recruits the kinase to Hsp90 for folding or modification. As the kinase is released from Hsp90, Cdc37^pS13^ will remain bound Hsp90 until dephosphorylated, leading to a unidirectional Hsp90-kinase cycle.

Agreeing with this hypothesis, the use of the longer CRaf^ED^ construct in our activity assays demonstrates the efficient dephosphorylation of both CRaf^pS338^ and CRaf^pS621^ by PP5. These experiments implicate PP5 in both CRaf^pS338^ dependent inactivation, but also in the dephosphorylation of CRaf’s “maturity marker”, CRaf^pS621^.^45^ Contrary to previous expectations, Cdc37^pS13^ was not dephosphorylated in the molybdate stabilized Hsp90^closed^:Cdc37:CRaf complex. Kinase steric inhibition, and Cdc37^pS13^ inaccessibility in the Hsp90:Cdc37:CRaf^KD^:PP5 structure support the lack of Cdc37^pS13^ dephosphorylation. The discrepancy between this and previous work might be explained by long experimental timeframes which might have led to complex dissociation.^32^ Meanwhile, Wandinger et al conducted experiments with CK2 phosphorylated Cdc37 which showed similar lack of Cdc37 dephosphorylation, potentially because of CK2 presence in their experiments.^26^

In our experiments, kinases sterically block the dephosphorylation of Cdc37 by PP5. This negates the hypothesis that PP5 can dephosphorylate Cdc37 in an Hsp90:Cdc37:kinase complex. Instead, we propose a model in which PP5 only dephosphorylates Cdc37 once the kinase has been released, thereby resetting Cdc37^pS13^ phosphorylation and preventing immediate kinase re-recruitment (Fig. 6b). This would bestow directionality to the Hsp90-kinase cycle and allow Hsp90 to release the kinase at a reset “basal” phosphorylation state.

In summary, Hsp90’s role in maintaining protein homeostasis goes beyond folding and activation to include client post-translational modification. Through the Hsp90:Cdc37:CRaf:PP5 cryo-EM structure, we further understand how Hsp90 activates PP5 and provides a scaffold for substrate dephosphorylation. Our biochemistry efforts suggest that PP5 may directly reset kinase dephosphorylation, while influencing Cdc37’s kinase recruitment to Hsp90. Further work will be required to provide a more complete mechanistic description of Cdc37^pS13^ dephosphorylation.

## Acknowledgements

We thank Agard Lab members for many helpful discussions; E. Nieweglowska for her mentorship and guidance; M. Tabios for his support and encouragement; C. Nowotny, D. Mozumdar and M. Moritz for help with Western blots; T. Kortemme and J. Gestwicki for their mentorship; C. Noddings, S. Pourmal, X. Liu, D. Asarnov for their help with CryoEM data processing; D. Bulkley, G. Gilbert, and Z. Yu from the W.M. Keck Foundation Advanced Microscopy Laboratory at the University of California, San Francisco (UCSF) for EM facility maintenance and help with data collection; M. Harrington and J. Baker-LePain for computational support with the UCSF Wynton Cluster. This work was supported in by NIH grant R35GM118099 (D.A.A) and NIH grants 1S10OD026881, 1S10OD020054, and 1S10OD021741 to the UCSF cryoEM facility.

## Author Contributions

M.J-G designed and executed protein preparation, biochemical experiments, cryo-EM sample preparation, data collection, data processing and model building. C.N. helped express yeast and mammalian constructs. D.C. helped conceive project idea and optimized yeast Hsp90:Cdc37:CRaf^KD^ expression and purification. F.W. optimized grids used for Cryo-EM, M.T. cloned PP5 mutants. D.A.A. edited, mentored, provided funding, and gave scientific advice throughout the project.

## Methods

### Data analysis and figure preparation

Figures were created using UCSF ChimeraX v.1.2.5 and Affinity Designer v1.10.5.^46,47^ Western blot data was analyzed using ImageJ 1.53k and plotted using Prism v.9.3.1 (GraphPad).^48^

### Individual component expression and purification

The Hsp90, Cdc37, and PP5 plasmids were transfected into E. coli BL21 cells, and then plated on antibiotic agarose plates. Overnight cultures were grown using Terrific Broth (TB), and then subsequently transferred to 2-6L of TB media. When O.D reached 0.6, cells were brought to 16°C for an hour after which they were induced with 1mM IPTG and the temperature was raised to 18°C, shaking overnight at 160RPM. The pellets were then spun down (4,500g for 10 min) and frozen until protein preparation.

The thawed pellets were then resuspended with Lysis buffer (500mM NaCl, 50mM Tris pH 7.8, 5% Imidazole, 5% Glycerol, 1 Protease Inhibitor Tablet/50mL, 0.5mM TCEP), and the sample was sonicated for five cycles, 1 minute/cycle, with a at least a minute between cycles, always submerged in ice. The sample was then spun down at 35k g for 30min at 4°C. The cellular lysate was then run through one to two HisTrap FF (5mL) Nickel columns at 5ml/min. The column was then loaded onto an Akta and washed with Wash Buffer (50mM Tris pH 7.8, 250mM NaCl, 0.5mM TCEP, 5% Glycerol, 30% Imidazole, PP5 buffer included 1mM Manganese), and then eluted with Elution Buffer (50mM Tris pH 7.8, 250mM NaCl, 0.5mM TCEP, 10% Glycerol, 400mM Imidazole, PP5 buffer included 1mM Manganese). This purified lysate was then concentrated down to 5-7mL with a Centricon tube and cleaved overnight (3C or TEV depending on plasmid).

The cleaved sample was then diluted down to ∼50mL with low salt buffer (20mM Tris pH 8.0, 1mM EDTA, 0.5mM TCEP, 5% Glycerol, PP5 buffer included 1mM Manganese), filtered, and loaded onto a 50mL superloop. This sample was then run through an Akta MonoQ 10/100 column Cytiva), with a gradient of low salt, and high salt buffers (Low Salt = no NaCl, High salt = 1M NaCl) and the peaks of interest were isolated and concentrated down to 250uL. The sample was then filtered, and loaded onto a Superdex 200 16/600 (Cytiva) column in storage buffer (20mM Hepes, 150mM KCl, 1mM EDTA, 1mM TCEP, 5% Glycerol). SDS Gel allowed for the inclusion of the cleanest protein fractions for further use. Samples were then frozen in liquid nitrogen and stored at -80°C.

BRaf expression and purification followed a similar protocol, with slight modifications: 0.75% IGEPAL was added to the lysis buffer, 20mM HEPES was used instead of 20mM Tris throughout the purification, and 10% glycerol was used until final size exclusion run.

### Complex expression and purification

Hsp90-Cdc37-CRaf complexes were purified from yeast and/or mammalian cells.

Yeast expression: Constructs were cloned into a 83 nu yeast expression vector, and transformed into JEL1 yeast strain using Zymo Research EZ transformation protocol and plated onto SD-His plates. After 3 days a colony was expanded into 200 mL overnight cultures. 1 L of YPGL media was inculcated with 10 mL of overnight culture. Cultures were induced with 2 % w/v galactose at an OD of 0.6 – 0.8 to induce expression. Temperature was reduced to 16°C and cultures were pelleted after 18 hours. Pellets were resuspended in minimal yeast resuspension buffer (20 mM HEPES-KOH pH 7.5, 150 mM KCl, 10% glycerol) and frozen drop wise into a container of liquid nitrogen.

Mammalian expression: Constructs were cloned into a pcDNA3.1 expression vector. Mammalian HEK293 cells were seeded at 0.5M/mL and allowed to reach a confluency of 3M/mL. Media was exchanged three hours before transfection and cells were allowed to recover. The Expi293TM Expression system kit and protocol was used for transformation. Cells were allowed to grow for 48 - 72 hours before they were spun down at 5k g, reconstituted with PBS and frozen drop by drop into liquid nitrogen.

The frozen cell drops were then cryomilled for 5 cycles (Precool 5 min, Run 1:30 min, Cool 2min, 10cps Rate). The samples were reconstituted with Strep binding buffer (20mM HEPES, 150mM KCl, 10mM MgCl, 1mM TCEP, 10% Glycerol, 0.05% Tween) with NaMo (20mM) added in cases of attempting to recover a more stable “closed” complex. The sample was loaded onto a StrepTrap (5mL) column at a rate of 5ml/min and consequently washed with 20mL of Strep binding buffer in an Akta at a flow rate of 5ml/min and then eluted with 10mL of Elution buffer (20mM HEPES, 150mM KCl, 10mM MgCl, 1mM TCEP, 10% Glycerol, 0.05% Tween, 10mM Desthiobiotin). The elution fractions were then concentrated down to 250uL and loaded onto the Superdex 200 16/600 (Cytiva) column with storage buffer (20mM Hepes, 150mM KCl, 1mM EDTA, 1mM TCEP, 5% Glycerol). The single peak was then concentrated, frozen with liquid nitrogen and stored at -80C.

### Dephosphorylation Assays

Reactions were carried out in reaction buffer (20mM Hepes, 50mM KCl, 10mM Mg2Cl, 1mM TCEP, 1mM EDTA) in PCR tubes. The Hsp90:Cdc37, or Hsp90:Cdc37:CRaf complexes were mixed at 4°C, and then allowed to reach 25°C on a thermocycler (∼2 min) before PP5 addition. PP5 was added, and the reaction begun upon mixing. Sample was removed from the thermocycler at each timepoint, and the reaction was quenched using SDS-DTT. Each reaction was repeated at least three distinct times with new sample. Dephosphorylation conditions were optimized such that the PP5 concentrations used were ideal for western blot visualization. 3uM of Cdc37^pS13^ and equimolar constituents were used for Cdc37 blots, with the final addition of 750nM of PP5^WT^ or PP5^mutant^. 1.5uM of Hsp90:Cdc37:CRaf complex with 75nM of PP5 were used for Fig. 1 experiments, while 3uM of Hsp90:Cdc37:CRaf complex and 150nM PP5 were used for PP5^mutant^ experiments.

The samples were run on Bolt™ 4-12% Bis-Tris gels (140V 65min) with the NEB# P7712 Color Protein Standard, Broad Range (10-250 kDa) for molecular weight differentiation. The gels were then transferred using Invitrogen’s iBlot® Gel Transfer Stacks Nitrocellulose, Regular following the transfer kit protocol (10 min transfer) and then stained with Ponzo stain for ∼5 min to ensure equal protein transfer and constant protein concentrations through timepoints (Supplementary files). The membranes were then incubated with 5% milk on a nutator for one hour at room temperature. Primary antibodies against Cdc37^pS13^ (1:5000 Phospho-CDC37 (Ser13) (D8P8F) Rabbit mAb #13248), CRaf^pS338^ (1:1000, # MA5-15176 Phospho-c-Raf (Ser338) Monoclonal Antibody(E.838.4)) or CRaf^pS621^(1:1000, #MA5-33196 Phospho-c-Raf (Ser621) Recombinant Rabbit Monoclonal Antibody) were then added to the membrane and nutated overnight at 4°C. The membrane was then washed with TBST (80mM Tris Base, 550mM NaCl, 1% Tween 20 (v/w)) three times, 15 min/wash. HRP Secondary antibody was then added to the membrane (1:10000, Anti-Rabbit NA9340V GE Healthcare UK Limited) and allowed to incubate on a nutator for 1h at RT. Next, the membrane was washed three times with TBST for 15 min/wash and exposed using the Thermofisher protocol and chemiluminescent agents (Pierce(tm) ECL Western Blotting Substrate, Catalog number: 32109).

An Azure biosystems imager was used to capture ponzo and chemiluminescence Western Blot images. The images were then analyzed using the ImageJ software.^48^ Each sample was run at least three separate times to ensure replicability. The western blots were then normalized by the zero timepoint (no PP5) using the Prism 9.3.1 (350) software. A one-phase linear decay curve was fit to dephosphorylation vs time data, and the rates of decay were compared using an ordinary one-way ANOVA test within Prism, corrected for multiple hypothesis testing using a Dunnett test. Multiple hypothesis testing was carried out within Prism. For visualization purposes, dephosphorylation rates for mutants were normalized to WT rates. All these values were then plotted with standard error of the mean error bars.

### Hsp90:Cdc37:CRaf:PP5 CryoEM Sample Preparation

Yeast purified Hsp90-Cdc37-CRaf^336-648^ complex was incubated with PP5 for 30 min at 4°C, then brought to room temperature, at which time 0.05% glutaraldehyde was added to the reaction and allowed to incubate for 15 min. The reaction was quenched with 50mM Tris buffer pH 8. The sample was then filtered (PVDF 0.1*µ*m), and 20*µ*L of sample was injected into the Ettan, where it ran through the Superdex 200 3.2/200 column (Cytiva) in running buffer (20mM Hepes pH 7.5, 50mM KCl, 1mM EDTA and 1mM TCEP). The fractions with Hsp90:Cdc37:CRaf:PP5 complex were separated from the PP5 excess, concentrated to ∼500nM and added to grids covered with a monolayer of graphene oxide derivatized with amino-PEG groups in a Vitrobot chamber (3uL of sample, 10°C, 100% humidity, 30s Wait Time, 3s Blot Time, -2 Blot Force) and plunged into liquid Ethane.^38,39^ Grids were stored in liquid nitrogen until use.

### CryoEM data acquisition and data processing

4160 micrographs were collected at 105000X magnification on a Titan Krios G3 (Thermo Fischer Scientific) with a Gatan K2 camera (0.835 Å/pix pixel size) at -0.8--1.8 µm defocus, 16 e-/pix*s, 0.025 sec/frame, per frame dose of 0.57e/A^2^*frame accumulating to a total dose of 69e^-^/Å^2^.

The images were motion corrected using UCSF Motioncor2, and their CTF estimated using CTFFIND-4.1.^49,50^ Only micrographs with a CTF fit <5Å were kept for Cryosparc blob picking. The picked particles were 2D classified (Cryosparc v.3.3.2) for the removal of high-resolution artifact particles, while the rest of the classes were kept for the next round of classification.^51^ The particles were then exported from Cryosparc and imported into Relion using csparc2star.^52^ The coordinates were used to re-extract particle picks using the Relion software (v3.1.3), where the rest of the data processing was carried out.^53^ We then carried out 3D classification, choosing all Hsp90 resembling classes. The approximately 1M particles were then refined and unbinned in a stepwise manner, until all particles were aligned onto Hsp90 at high resolution. An Hsp90:Cdc37:kinase^C-lobe^ model was used for refinement to side-step orientation confusion (low pass filtered from PDB: 5fwk). Once aligned on Hsp90, regions of interest were subtracted, and focused classified without alignment. Classes of interest were un-subtracted and refined until no improvement was seen. Subsequently, cycles of post-processing, per particle CTF refinement and refinement were repeated until no there was no resolution improvement. Finally, these maps were sharpened in Phenix using local anisotropic sharpening, and these final maps were used for model fitting.^54^

### Model Building and refinement

Hsp90:Cdc37:kinase Cryo-EM structures (PDB: 5fwk and unpublished D. Coutandin) and PP5 crystal structures (PDB: 1wao, 5hpe) were used for model building. The PP5 linker was built using RosettaCM after whole domain docking of both PP5 domains.^55,56^ To ensure best model fit, the PP5 linker was built N to C terminally, and C to N terminally to ensure convergence. The Cdc37^MD^ model from 5fwk was improved by using RosettaCM and ISOLDE.^57^ PhenixRefine was used after model docking/building to correct complex geometries, and RosettaRelax was used to consider protein chemistry in lower resolution parts of the model. ISOLDE was finally used to improve clashes, Ramachandran outliers, and rotamer fits.

