## Supplementary Figures for "Hsp90 provides a platform for kinase dephosphorylation by PP5"

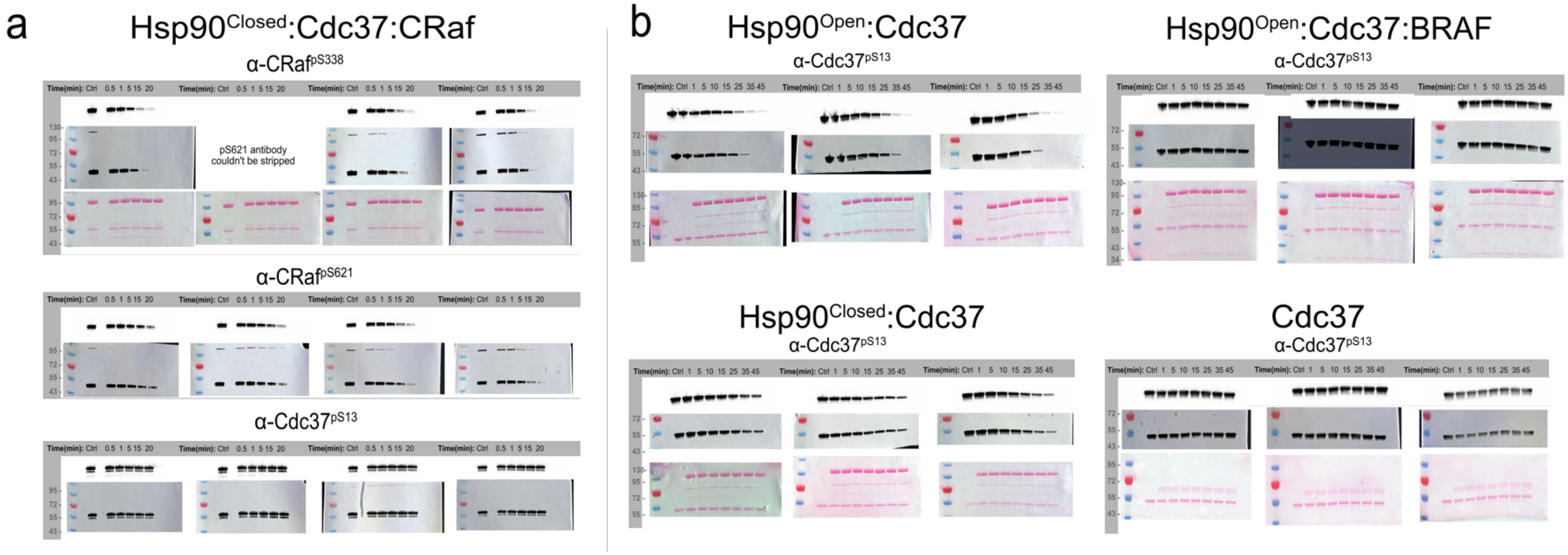


**Supplementary Fig. 1: Western blot replicate data. a** Purified CRaf-complex(1.5uM) was incubated with PP5 (75nM) at 25°C, ran on a gel transferred to a nitrocellulose membrane, ponzo stained and blotted for α-CRaf$^{pS338}$, α-CRaf$^{pS621}$, and α-Cdc37$^{pS13}$. **b** Reconstituted complex (3uM) was incubated with PP5 (750nM) at 25°C, ran on a gel, transferred to a nitrocellulose membrane, ponzo stained and blotted for α-Cdc37$^{pS13}$. CRaf dephosphorylation happens at a faster rate, so PP5 concentration was lowered, and time increments shortened. Equal amounts of protein can be seen in ponzo stained membrane throughout experimental timepoints.

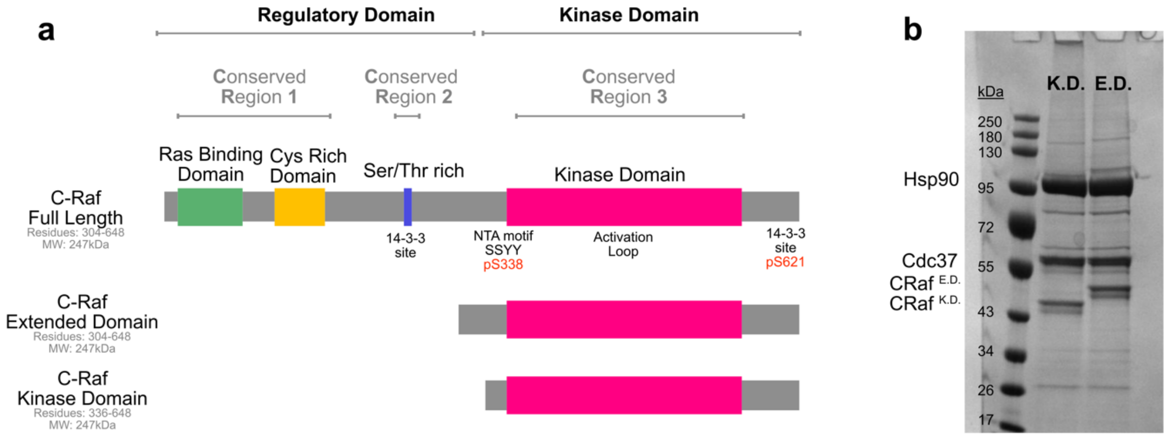


**Supplementary Fig. 2: CRaf topology and constructs. a** CRaf consists of a Regulatory and Kinase domain. The CRaf regulatory domain contains a Ras Binding Domain, a Cysteine Rich Domain, and important phosphorylation sites such as pS259 which binds 14-3-3 chaperones.The CRaf kinase domain consists of two lobes, the beta sheet heavy N-lobe and alpha helical C-lobe. ATP binds between these two lobes. Important phosphorylation sites around the kinase domain are highlighted in red and consist of the N-terminal pS338 and the C-terminal pS621. Two shorter CRaf constructs were used to increase solubility of CRaf and enable further biochemistry. The Extended domain (CRaf$^{ED}$ ) was used for all biochemical experiments, while the Kinase domain (CRaf$^{KD}$) was used for structural characterization. **b** Hsp90:Cdc37:CRaf complexes were purified from ExpiHEK293 cells with coexpressed Hsp90 and Cdc37.

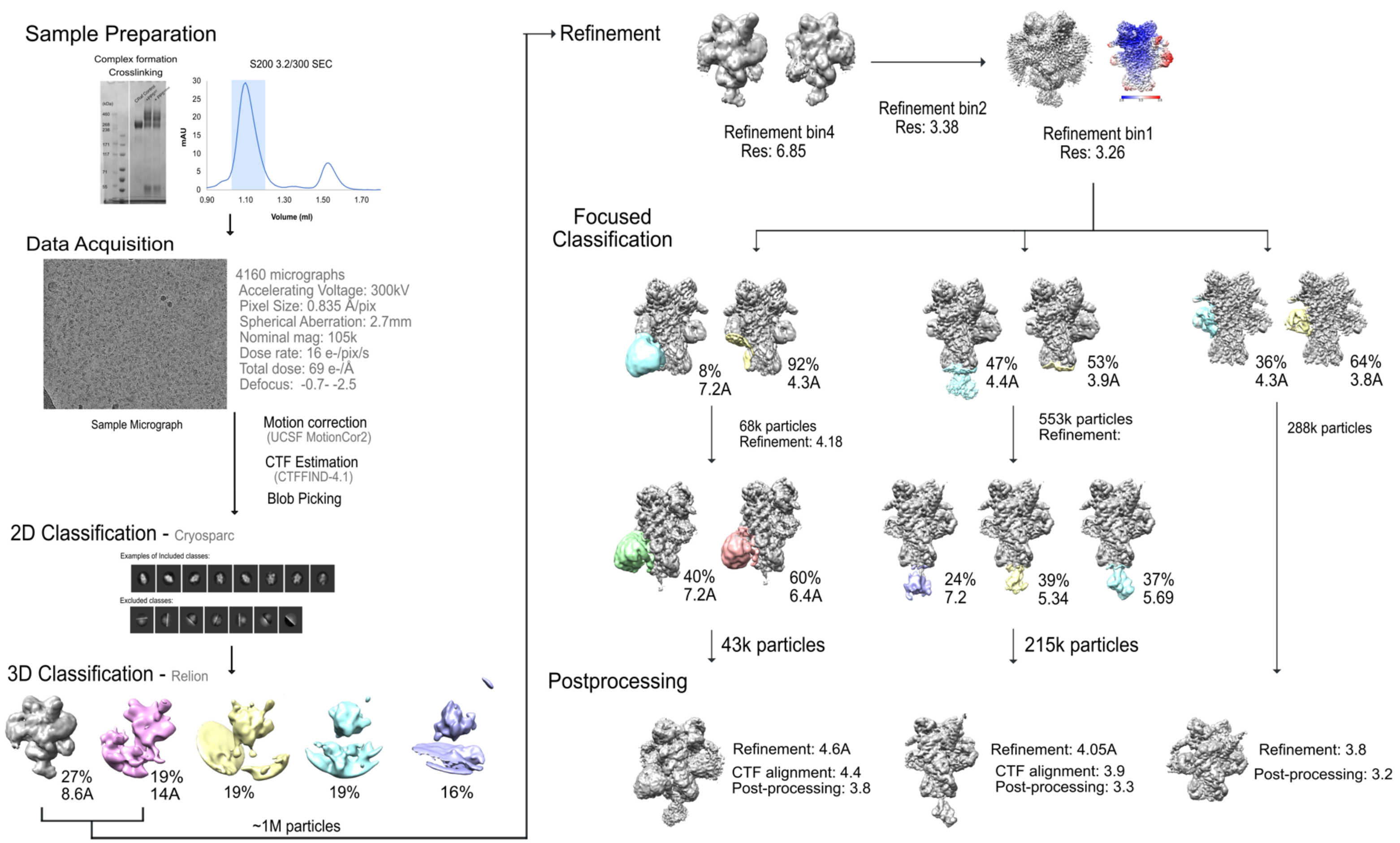


**Supplementary Fig. 3:** Sample preparation and cryo-EM data processing Human Hsp90$^{Closed}$-Cdc37-CRaf complex was purified from yeast, and PP5$^{H304A}$ was purified from E. Coli. Hsp90 complex (2uM) was mixed with PP5$^{H304A}$ (6uM) for 30 min on ice, and then brought to RT where Gluteraldehyde (0.05%) was added and allowed to crosslink for 15min until quenching with 50mM Tris pH 7.8. The crosslinked complex was then run over an S200 column (3.2/300) to remove excess PP5. The fractions from the major peak (~1.1mL) were collected and concentrated 3 fold. This sample wasfrozen on PEG Amino functionalized Gold carbon quantifoil grids using a Vitrobot machine (10C, 100% humidity, 30s Wait Time, 3s Blot Time, -2 Blot Force).  4160 micrographs were acquired using a Krios microscope (105kX mag). The images were then motion corrected, and their CTF estimated. Only micrographs with a CTF fit <5A were kept for Cryosparc blob picking. The particles were then 2D classified to remove high resolution artifact particles and ensure sample quality. The particle [icks from the chosen 2D classes were then imported into Relion (using csparc2star by D. Asarnov) and 3D classification begun. One round of classification led to ~1M Hsp90 like particles, which were then refined and unbinned. From various rounds of focused classification and consequent Post-processing led to the final maps used to model fitting and composite map creation.

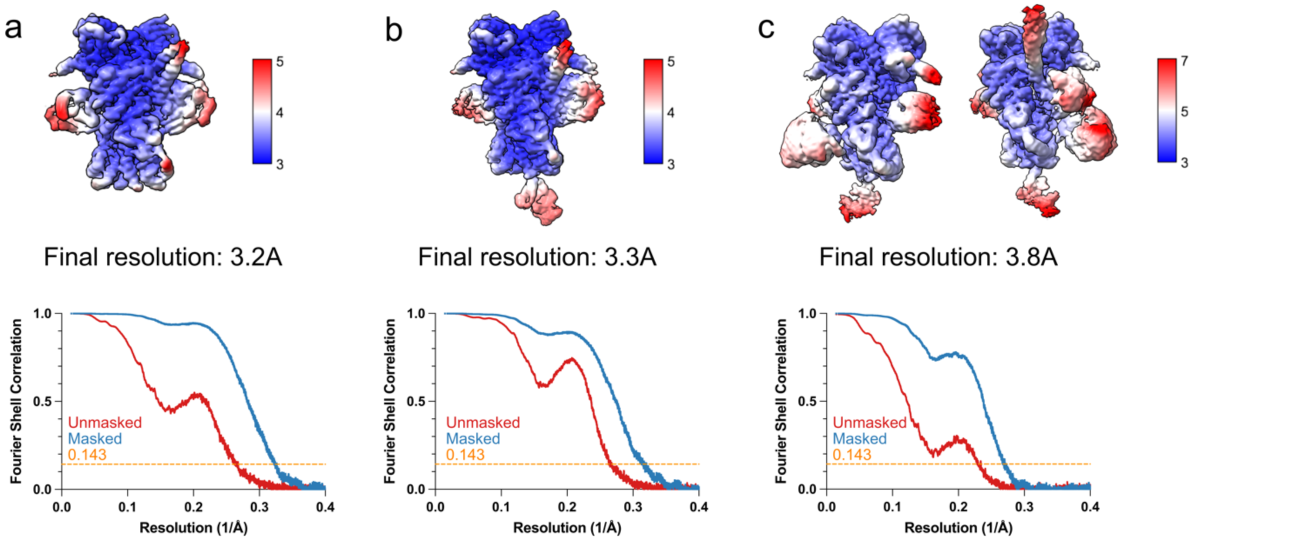


**Supplementary Fig. 4**: Three final maps were used for composite map formation. All maps showed the highest resolution at Hsp90, with lower resolution in the other domains. a Focused classification of the CRaf N-lobe, **b** focused classification of the PP5 TPR domain shows higher resolution where the $\alpha$helix of PP5 contacts Hsp90, and **c** focused classification of the PP5 catalytic domain shows higher resolution at the $Hsp90^{MD}$-PP5 contact points.

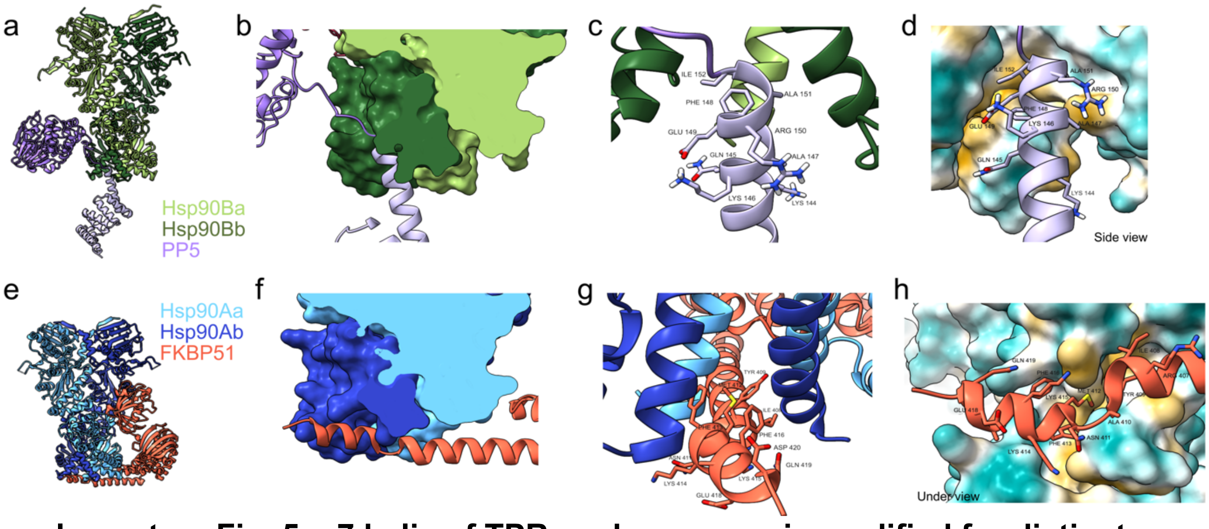


**Supplementary Fig. 5: a7 helix of TPR cochaperones is modified for distinct Hsp90 interactions**. **a,e** Hsp90 interacts with PP5 and FKBP51 via C-terminal interactions. **b,f** The cochaperone's a7 helices interact with the Hsp90 C-terminal groove at an almost ~90° angle difference from each other. **c,g** While hydrophobic residues line one face of the FKBP51 helix, the PP5 $\alpha$7 helix is polar at only two residues, allowing the rest of its helix to make polar interactions with the Hsp90 side.

**d,h** The PP5 and Fkbp51 $\alpha 7$ helices interact distinctly with the Hsp90$^{CTD}$ groove hydrophobic pocket (blue hydrophilic, yellow hydrophobic).

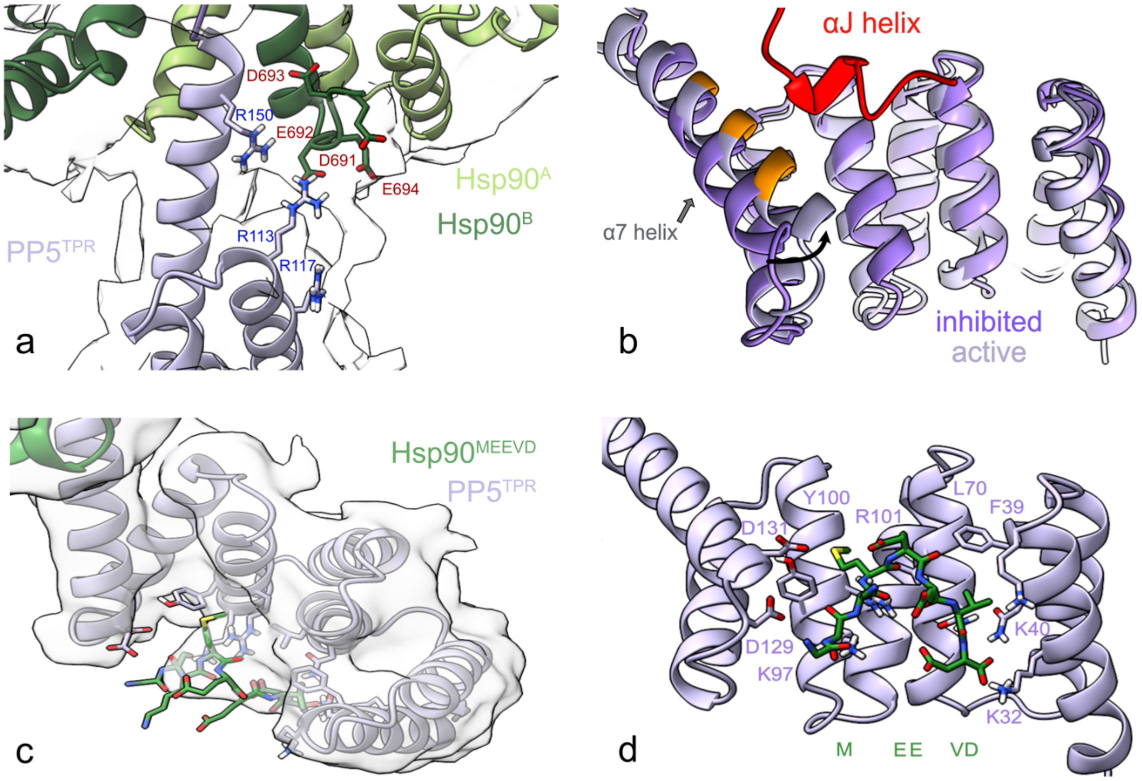


**Supplementary Fig. 6: PP5's TPR domain interacts with the Hsp90 tail and MEEVD motif, but not with PP5's $\alpha$J helix upon activation**. **a** Density can be seen reaching from the Hsp90$^{CTD}$ towards PP5's basic residues. **b** Overlay of PP5's TPR domain in an inhibited (PDB: 1wao) and active state shows the clashing (orange) of the PP5 $\alpha$J helix with the active PP5 $\alpha 7$ helix. **c** Density can be seen for Hsp90's MEEVD peptide as it interacts with PP5's basic patch. **d** A homologous structure of a TPR-MEEVD interaction was docked to highlight the location of key MEEVD binding residues on PP5. (PDB: 6q3q).

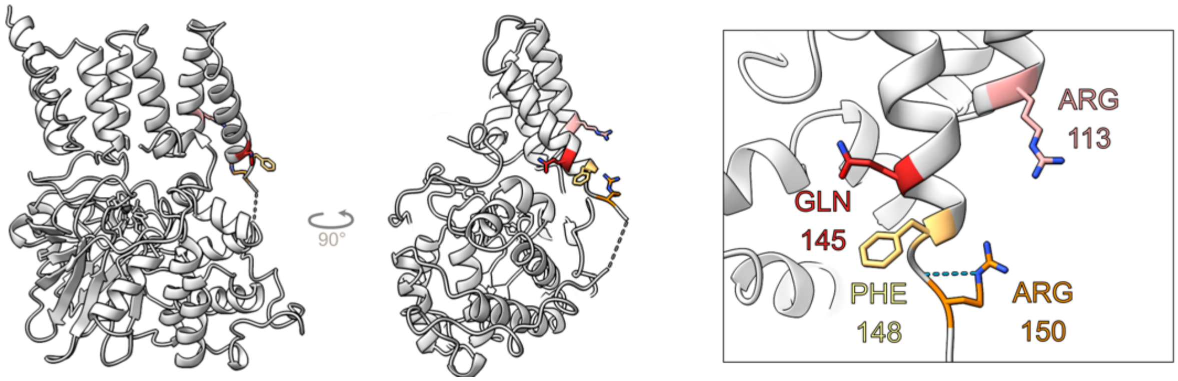


**Supplementary Fig. 7: TPR mutants surface exposed in inhibited PP5 structure.** PP5's TPR mutants face away from PP5's catalytic site making no salt bridges with other residues and sitting > 8Å away from the catalytic domain. The $\alpha$7 helix on the TPR domain will be flipped and bound to Hsp90's C-terminal domains.

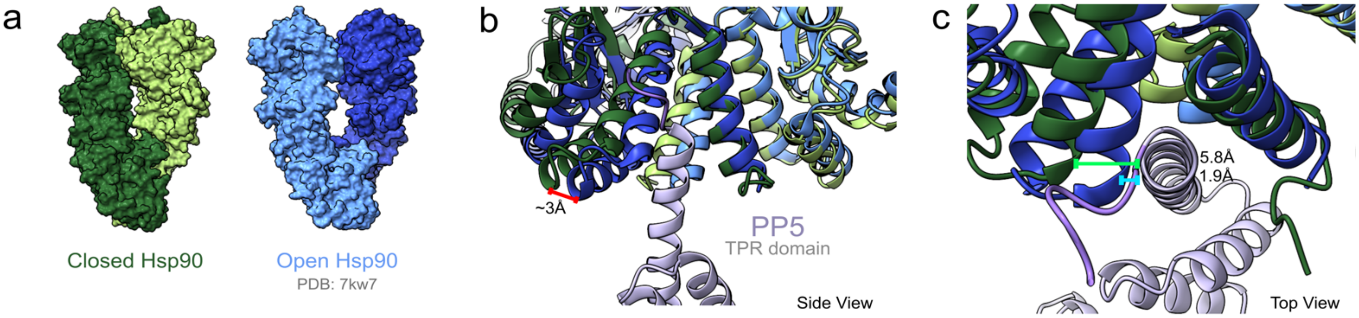


**Supplementary Fig. 8: Open Hsp90 might affect PP5 binding to Hsp90. a** Closed Hsp90 dimer as seen in our current work is contrasted with the loading complex semi-open Hsp90 dimer (PDB: 7kw7). **b** A side view of an overlay of the Hsp90 open and closed models shows a shift in the Hsp90$^{CTD}$ which narrows the Hsp90$^{CTD}$ groove used for PP5 binding by approximately 3Å. **c** A top down view of the overlayed Hsp90 models shows a distance of less than 2Å between the PP5 $\alpha$7 helix and the Hsp90$^{CTD}$ in a semi-open state. This would lead to a probable clash between Hsp90 and PP5 unless the PP5 $\alpha$7 helix moves out of the Hsp90$^{CTD}$ groove or interacts with the opposite Hsp90 protomer more closely.

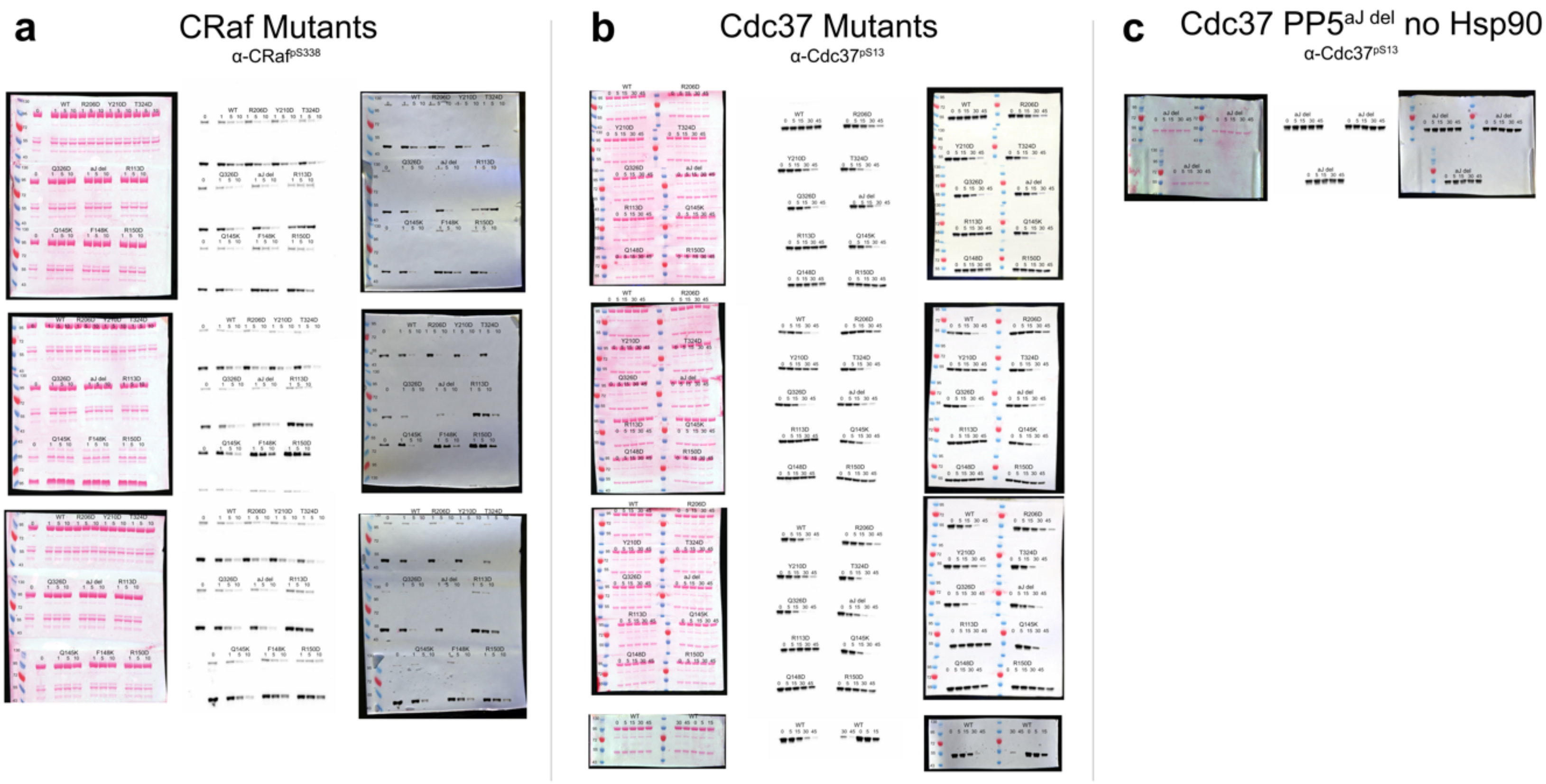


**Supplementary Fig. 9: Mutant western blot replicate data.** PP5 mutants were tested for rates of dephosphorylation as described in Supplementary Fig. 1. **a** Mammalian purified $Hsp90^{Closed}$:Cdc37:$CRaf^{ED}$ (3uM) was incubated with PP5 mutant (150nM). **b** E coli purified, CK2 phosphorylated $Cdc37^{pS13}$ (3uM) was incubated with Hsp90 and PP5 mutants (750nM). Additional mutants not considered in the manuscript for lack of clarity are included here. The Wild type (WT) condition was repeated twice more. **c** Cdc37 (3uM) was incubated with $PP5^{aJ\ helix\ del}$ (750nM) in the absence of Hsp90.

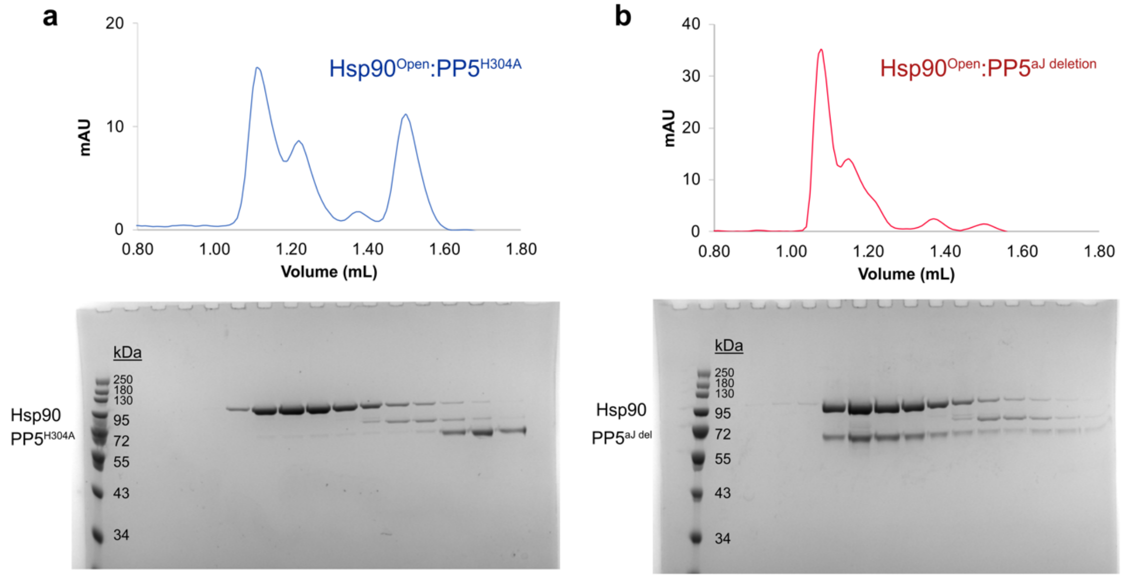


**Supplementary Fig. 10: αJ helix Truncation leads to increased Hsp90:PP5 complex coelution.** Hsp90 dimer was incubated with either catalytically dead PP5$^{H304A}$ or PP5$^{aJ\ del}$ at 4°C for 30min before loading the sample onto the S200 3.2/300 (Cytiva) column in SEC buffer (20mM HEPES, 50mM KCl, 10m $MgCl_{2,}$ 1mM EDTA, 1mM TCEP) and allowed to flow through at a flow rate of 0.04ml/min. 50uL aliquots were eluted and ran on SDS gels for component visualization. **a** PP5$^{H304A}$ barely coelutes with Hsp90, while **b** PP5$^{aJ\ del}$ strongly coelutes with Hsp90.
